# MIC13 and SLP2 seed the assembly of MIC60-subcomplex to facilitate crista junction formation

**DOI:** 10.1101/2023.09.04.556207

**Authors:** Ritam Naha, Rebecca Strohm, Jennifer Urbach, Ilka Wittig, Andreas S. Reichert, Arun Kumar Kondadi, Ruchika Anand

## Abstract

The MICOS complex subunit MIC13 is essential for mitochondrial cristae organization. Mutations in *MIC13* cause severe mitochondrial hepato-encephalopathy displaying defective cristae morphology and loss of the MIC10-subcomplex. Here we identified stomatin-like protein 2 (SLP2) as an interacting partner of MIC13 and decipher a critical role of SLP2 as an auxiliary MICOS subunit, modulating cristae morphology. SLP2 provides a large interaction hub for MICOS subunits and loss of SLP2 leads to drastic alterations in cristae morphology. Double deletion of SLP2 and MIC13 showed reduced assembly of core MICOS subunit, MIC60 into MICOS and dispersion of MIC60-specific puncta, demonstrating a critical role of SLP2-MIC13 in MICOS assembly and crista junction (CJ) formation. We further identified that the mitochondrial i-AAA protease YME1L in coordination either with MIC13 or SLP2 differentially regulates MICOS assembly pathways thereby interlinking MIC13-specific or scaffolding-specific role of SLP2 with quality control and assembly of the MICOS complex. YME1L- depletion in *MIC13* KO could restore MIC10-subcomplex and reform the nascent CJ. Taken together, we propose ‘seeder’ model for MICOS assembly and CJ formation, where SLP2- MIC13 seed the assembly of MIC60 into MICOS complex and promote the formation of CJ by regulating the quality and stability of MIC10-subcomplex.

## Introduction

Mitochondria with their multifaceted roles are involved in many cellular functions, in addition to energy conversion, namely calcium signalling, lipid metabolism, ROS production, amino acid metabolism, iron-sulfur cluster synthesis and regulation of apoptosis (Monzel, Enríquez et al., 2023). The inner membrane (IM) of mitochondria is versatile and adapts according to the bioenergetic demands of the cell. Cristae are infoldings of the IM that provide the characteristic wrinkled shape of IM and offer large surface area for housing electron transport chain (ETC) complexes. The intricate three-dimensional (3D) arrangement of the IM was better illustrated upon use of electron tomography techniques in 1990s leading to the proposal of crista junction (CJ) model. CJs are small openings at the neck of individual cristae with a high inward-directed curvature (Perkins, Renken et al., 1997). Due to their small diameter, CJs were proposed as a diffusion barrier (Mannella, Lederer et al., 2013, Perkins et al., 1997) subdividing mitochondria into various subcompartments and thus modulating several mitochondrial functions. CJ could modulate lipid transfer and metabolite exchange in the mitochondria. The remodelling of CJ during apoptosis lead to release of cytochrome *c* and initiation of apoptotic cascade (Scorrano, Ashiya et al., 2002). However, how these intricate structures of cristae and CJ with steep membrane curvatures are formed and remodelled remain elusive for decades. Several recent findings provide insights on this very significant yet technically challenging question. MICOS ‘mitochondrial contact site and cristae organising system’ complex has emerged as a critical player in formation of cristae and CJ (Anand, Reichert et al., 2021, Mukherjee, Ghosh et al., 2021, Stephan, Brüser et al., 2020). Live-cell super-resolution nanoscopy showed that cristae and CJs are highly dynamic and undergo cycles of fusion and fission at a timescale of seconds in a MICOS-dependent manner (Kondadi, Anand et al., 2020). Mammalian MICOS complex contains seven subunits that are further divided into two subcomplexes, the MIC60- subcomplex (MIC60-MIC19-MIC25) and the MIC10-subcomplex (MIC10-MIC13-MIC26- MIC27). Deletion of individual subunits of the MICOS complex causes aberrant cristae morphology and accumulation of cristae stacks or concentric rings to a variable degree (Kondadi et al., 2020, Stephan et al., 2020). MIC60 and MIC10 are the most important subunits and are shown to harbour membrane bending activity (Barbot, Jans et al., 2015, Bock-Bierbaum, Funck et al., 2022, Bohnert, Zerbes et al., 2015, Hessenberger, Zerbes et al., 2017, Tarasenko, Barbot et al., 2017). MIC19 and MIC25 are coiled-coil helix coiled-coil helix (CHCH) family proteins and important to assist MIC60 in the formation of CJs (An, Shi et al., 2012, Sakowska, Jans et al., 2015). MIC13 is a small protein which is proposed as an essential component for maintaining contact between the two MICOS subcomplexes (Anand, Strecker et al., 2016, Guarani, McNeill et al., 2015, Urbach, Kondadi et al., 2021, Zerbes, van der Klei et al., 2012). MIC26 and MIC27 are homologous proteins that belong to the apolipoprotein family and required for cristae morphology, integrity of respiratory chain complexes and cardiolipin levels (Anand, Kondadi et al., 2020). MIC27 has the ability to bind cardiolipin (Weber, Koob et al., 2013). The interaction of the MICOS subunits with outer membrane (OM) proteins SAMM50, metaxin1 (MTX1), metaxin2 (MTX2) and DNAJC11 forms a larger ‘mitochondrial intermembrane space bridging’ (MIB) complex that is responsible for formation of contacts between IM and OM (Huynen, Muhlmeister et al., 2016, Tang, Zhang et al., 2020, Xie, Marusich et al., 2007). Mutations in several MICOS subunits have been linked to various human diseases but the pathobiology of mitochondrial diseases that are occurring due to the non-ETC genes are not studied in detail. Mitochondrial diseases account of a large class of inborn errors of metabolism with the prevalence rate of 1.6 in 5000 (Stenton & Prokisch, 2020). Yet, no curative treatment is known. Mutations of *MIC60, MIC26 and MIC13* have been found in several severe human diseases (Beninca, Zanette et al., 2021a, Guarani, Jardel et al., 2016, Peifer-Weiß, Kurban et al., 2023, Russell, Whaley et al., 2019, Tsai, Lin et al., 2018, Zeharia, Friedman et al., 2016). The question remains how cristae and CJs are formed and maintained during the healthy or pathological conditions and how does specific MICOS subunit contribute to these processes. In this study, we specifically focused on determining the molecular role of MIC13 in cristae formation that could provide novel insights into the relevant pathobiology.

MIC13 is not well characterized and no homology was found with other protein families or domains (Urbach et al., 2021). Loss of MIC13 causes concomitant total loss of MIC10, MIC26 and partial loss of MIC27, making it difficult to assign the specific phenotype to any of the proteins involved (Anand et al., 2016, Guarani et al., 2015). We had reported two conserved motifs, the ‘GxxxG’ and the ‘WN’ motif, in MIC13 that were important for its stability and function (Urbach et al., 2021). Nevertheless, it is an important component of the MICOS complex and mutations in *MIC13* are associated with severe form of mitochondrial hepato-encephalopathy in children (Guarani et al., 2016, Russell et al., 2019, Zeharia et al., 2016). The patients die at very early age ranging from a few months to 5 years. The pathology included multi-system failure in brain, liver, kidney and heart (Godiker, Gruneberg et al., 2018, Guarani et al., 2016, Russell et al., 2019, Zeharia et al., 2016). Neurological defects included cerebellar and optic atrophy, acquired microcephaly and hypotonia. Most patients also showed clear signs of liver disease accompanied by acute liver failure. Increased plasma levels of lactic acid, methionine, tyrosine and kerbs cycle intermediates and increased excretion of 3-methylglutaconic acid was reported. In all the cases documented, MIC13 levels were not detectable, indicating the complete loss of MIC13 in these pathologies.

To identify the molecular role of MIC13, we set out to identify its interaction partners. Using mass-spectrometry (MS) coupled with coimmunoprecipitation (co-ip) of MIC13, we detected Stomatin-like protein 2 (SLP2) as one of the highly enriched proteins in MIC13 interactome. A link to MICOS regulation or cristae morphogenesis has not been reported for SLP2. Here, we identified SLP2 as an auxiliary subunit of MICOS. SLP2 provides a scaffold to form a large interaction hub for all known MICOS subunits. Our results show a novel multi-layered role of SLP2 in regulating MICOS assembly and cristae morphogenesis. SLP2 was specifically required for the stability of MIC26 and its incorporation in the MICOS complex by regulating YME1L-mediated MIC26 proteolysis. Moreover, the combined depletion of MIC13 and SLP2 accentuates defects in MIC60 assembly, emphasizing their collaborative roles in modulating assembly kinetics and formation of MIC60 puncta. Next to novel roles of SLP2 and YME1L, our study elucidates MIC13-specific role in MICOS assembly and cristae morphogenesis, which is important for understanding the MIC13-associated pathophysiology. We further introduce a ‘seeder model’ of MICOS assembly, wherein SLP2, along with an assembled MIC10-subcomplex (‘seeder’ complex), facilitates the efficient incorporation ‘seeding’ of MIC60 into the holo-MICOS-MIB complex, ensuring mitochondrial integrity and formation of CJ and contact between IM and OM.

## Results

### Determining the MIC13 interactome

To unravel the unidentified function of MIC13 in regulating MICOS and/or processes independent of MICOS, we determined the MIC13 interactome. Isolated mitochondria from Flp-In T-REx HEK293 cells were subjected to co-ip using agarose beads conjugated with MIC13 antibody and the eluate fraction was analysed by MS to identify the proteins which were specifically and significantly enriched in wild type cells compared to *MIC13* knockout (KO) cells. The analysis led to identification of numerous proteins which constituted the interactome of MIC13 in mammalian cells (Fig 1A, Supplementary Table 1). Many of the identified proteins belonged to MICOS and MIB complex or their known interactors, which demonstrates the specificity of the results and highlights the central role of MIC13 in MICOS-MIB regulation. We also found SLP2 as a novel interaction partner which showed the highest fold enrichment in significance upon statistical analysis (Fig 1A). SLP2 belongs to the SPFH (stomatin, prohibitin, flotillin, HflC/K) superfamily of scaffolding proteins that can form microdomains in the membrane by local lipid-protein interactions. SLP2 is an IM protein which can bind cardiolipin (Christie, Lemke et al., 2011) and regulate many mitochondrial functions including biogenesis, proteolysis and morphology during stress-induced mitochondrial hyperfusion (SIMH) (Tondera, Grandemange et al., 2009). A direct role of SLP2 in regulation of MICOS-MIB and cristae morphology has not yet been reported, prompting us to study this possibility in more detail.

**Figure 1.**
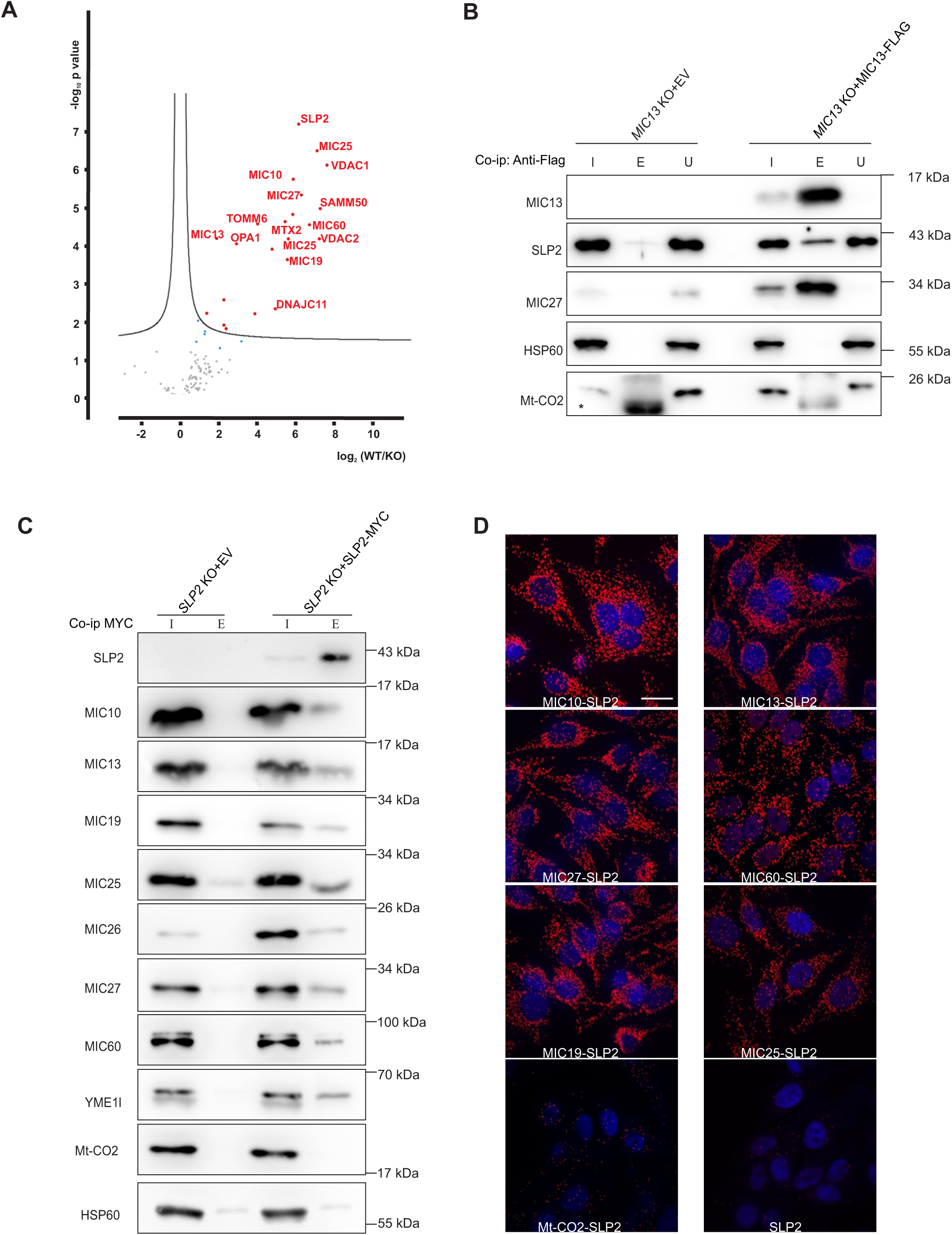
SLP2 is identified as a novel MIC13 interacting partner. **A**, Interactome of MIC13 with co-ip (co-immunoprecipitation) coupled mass spectrometry revealed SLP2 as a novel interactor of MIC13. **B,** The interaction between SLP2 and MIC13 was validated by co-ip using FLAG antibody in isolated mitochondria from *MIC13* KO cells stably expressing MIC13-FLAG or empty vector (EV) pMSCVpuro as background control. I: input lanes represent loading of 10% of total lysates, E: eluate represent proteins eluted from anti-Flag M2 beads, U: unbound fraction. * non-specific IgG bands. **C,** Co-ip probed for SLP2-MICOS interaction with isolated mitochondria from *SLP2* KO stably expressing pMSCVpuro EV (background control) or SLP2- MYC. Co-ip was performed using MYC-Trap agarose beads. I: Input fraction (10% of total lysate), E: Eluate fraction. YME1L was used as a positive interactor of SLP2 whereas Mt-CO2 and HSP60 served as non-interactors. All the MICOS subunits were present in the elution fraction from SLP2-MYC co-ip. **D,** Proximity ligation assay (PLA) in HeLa cells with antibodies against MICOS subunits and SLP2. PLA signals are shown as red spots indicating respective protein interactions. SLP2 alone and Mt-CO2 & SLP2 antibodies were probed as negative controls.

### SLP2 stably interacts with all MICOS subunits forming an interaction hub

In order to validate the interaction between MIC13 and SLP2 (Fig 1A), the elution fraction from the MIC13-FLAG co-ip was subjected to western blot (WBs) analysis and probed with an SLP2 antibody. The SLP2 band intensity was substantially higher in *MIC13* KO cells expressing MIC13-FLAG compared to *MIC13* KO expressing an empty vector (EV) confirming the interaction of MIC13 with SLP2 (Fig 1B). The absence of matrix protein HSP60 and ETC protein Mt-CO2 in the elution fraction showed the specificity of the co-ip experiment (Fig 1B). Of note, only a fraction of SLP2 interacted with MIC13 as other SLP2 remained in unbound fraction compared to the positive interactor MIC27. Further, we tested the MIC13-SLP2 interaction using SLP2 as a bait in a co-ip experiment. For this, we generated *SLP2* KO cells using CRISPR-Cas9 system and stably expressed SLP2 with a MYC tag at its C-terminus. Co-ip was performed using MYC-trap agarose and the elution fraction was probed for antibodies against SLP2, YME1L (known SLP2 interactor), all MICOS subunits, HSP60 and Mt-CO2 (as negative controls). The presence of SLP2 and YME1L and the absence of HSP60 and Mt-CO2 in the elution fraction showed the specificity of the co-ip experiment (Fig 1C). All MICOS subunits, and not only MIC13, were present in the elution fraction, showing that SLP2 can directly or indirectly interact with all the MICOS subunits (Fig 1C). Proximity ligation assay (PLA) can be used to determine and visualize the proximity (interaction) between two proteins of interest. PLA using antibodies specific to SLP2 and individual MICOS subunits showed numerous punctae in each cell which mark the interaction between SLP2 and individual MICOS subunits compared to negative controls of Mt-CO2 & SLP2 or only SLP2 antibody (Fig 1D). Overall, several lines of evidence confirm the specific and reciprocal interaction of SLP2 with the MICOS complex.

Due to the known limitations of co-ip experiments, we cannot specify whether SLP2 interacts individually with each MICOS subunit or whether there is a hierarchy in the interaction of MICOS subunits with SLP2. To determine if the interaction between SLP2 and MICOS subunits relies on any particular MICOS subunit, we decided to perform co-ip experiments in cells deleted for individual MICOS subunit. We generated KO cells lacking individual MICOS subunits (MIC10, MIC26, MIC27, MIC19, MIC25 and MIC60) in HEK293 cell lines and stably expressed SLP2-Myc in these cell lines. As expected, the *MIC10* KO and *MIC13* KO cells showed loss (or decrease) of all subunits of the MIC10-subcomplex (MIC10/13/26/27) while KO of either *MIC26* or *MIC27* showed no drastic change in other MICOS subunits (Fig 2A). Among the subunits of MIC60-subcomplex, *MIC60* KO cells showed a drastic decrease in steady state levels of all other MICOS subunits, whereas *MIC19* KO cells showed reduced MIC10, MIC60 and MIC13 (Fig 2B). *MIC25* KO cells showed no drastic defect in the levels of other MICOS subunits (Fig 2B). Despite the known and observed loss of subunits of the MIC10-subcomplex in *MIC10* KO and *MIC13* KO cells, we clearly observed that SLP2 still interacts with YME1L, MIC19, MIC25 and MIC60 (Fig 2A). Similarly, SLP2 still interacted with the remaining MICOS subunits in *MIC26* KO, *MIC27* KO and *MIC25* KO cells (Fig 2A, 2B). As *MIC60* KO cells showed very low levels of all MICOS subunits in the input fraction, we included an over-exposed blot in addition at the right (Fig 2B). This clearly showed that the remaining MIC13, MIC26, MIC27 and MIC19 subunits in *MIC60* KO could still interact with SLP2 (Fig 2B). Altogether, we conclude that SLP2 can stably interact with any remaining MICOS subunits even when other individual MICOS subunits are lost.

**Figure 2.**
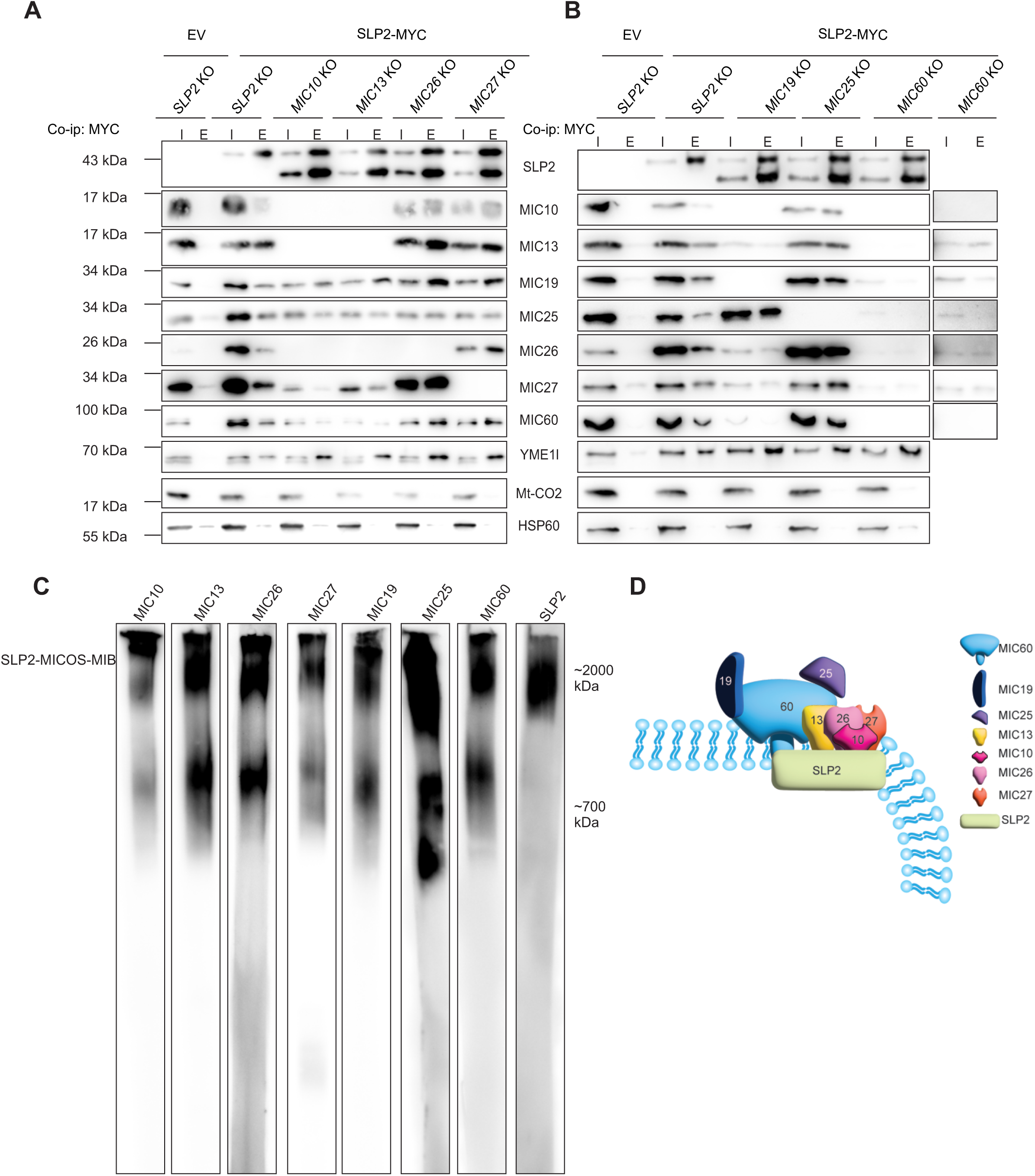
SLP2 forms an interaction hub with MICOS complex proteins. **A**, Co-ip-western blot analysis from *SLP2* KO or MIC10-subcomplex KO (left) and **B**, *SLP2* KO or MIC60- subcomplex KO (right) stably expressing pMSCVpuro EV or SLP2-MYC showed that SLP2 can stably interact with any remaining MICOS subunits even upon the loss of individual MICOS subunits. Due to low abundance of MICOS proteins in *MIC60* KO cells, overexposed blots were represented showing independent interaction of MIC13, MIC26, MIC27 and MIC19 with SLP2 in absence of MIC60. I: Input fraction (10% of total lysate), E: Eluate fraction. **C,** BN- PAGE with isolated mitochondria from WT cells revealed a co-migration pattern of SLP2 with higher molecular weight MICOS complex. **D,** Scaffolding model depicting interaction of SLP2 as an auxiliary MICOS subunits shows that SLP2 provides a scaffold for interaction of MICOS subunits.

We next performed a blue-native gel electrophoresis (BN-PAGE) to determine the high-molecular weight complexes of SLP2 and MICOS. Normally, MICOS subunits are distributed in two large complexes; a higher MICOS complex with around 2000 kDa size that is shown to also include MIB subunits and the lower molecular weight MICOS complex with the size around 700 KDa (Anand et al., 2016). We found that SLP2 forms a high molecular weight complex which runs parallel to the high-molecular weight MICOS complex at around 2000 kDa (Fig 2C). This observation was also verified in the complexome profiling data from HEK293 cell (Anand et al., 2016) which shows the co-clustering of SLP2 and MICOS subunits at the region of 2000 kDa (Supplementary Fig 1). Overall, these results show that SLP2 provides a scaffold to form an interaction hub with the individual MICOS complex in the IM of mitochondria (Fig 2D). Therefore, we suggest that SLP2 is an auxiliary subunit of MICOS complex.

### SLP2 and YME1L determine the stability of MIC26

Next, we investigated the significance of the SLP2-MICOS interaction and asked whether loss of SLP2 affects the steady state levels of MICOS subunits or the MICOS complex. Using WBs, we analysed the levels of MICOS subunits in *SLP2* KO cells and found a drastic reduction in the steady state levels of MIC26, while the steady state of other MICOS subunits were largely unaltered (Fig 3A). The MICOS assembly using a BN-PAGE showed that MIC26 was sparsely present in the MICOS complex as expected from the steady state levels, however the incorporation of most other MICOS subunits into the MICOS complex appeared comparable to WT (Fig 3B).

**Figure 3.**
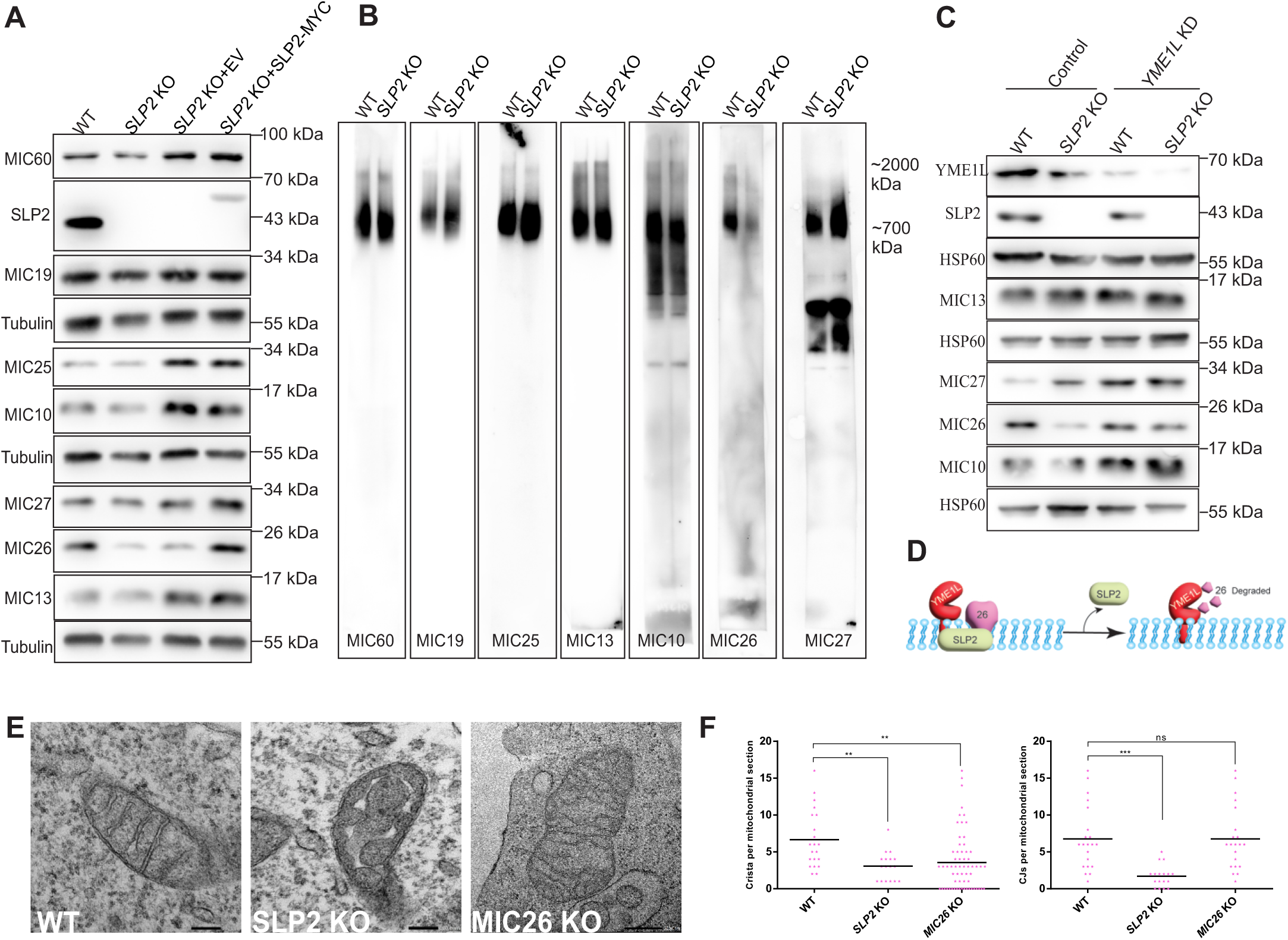
Loss of SLP2 leads to aberrant cristae structure and reduced MIC26 levels. **A**, Steady state levels of MICOS proteins with western blot analysis from WT, *SLP2* KO and *SLP2* KO cells stably expressing pMSCVpuro EV or SLP2-MYC. **B,** BN-PAGE of isolated mitochondria from WT and *SLP2* KO cells stained for MICOS subunits. **C,** Western blot analysis of steady state levels of MICOS proteins from WT and *SLP2* KO cells stably expressing pGIPZ-control shRNA or YME1L shRNA (knockdown represented as KD). **D,** A model depicting the role of SLP2 in stabilizing MIC26 by regulating YME1L-mediated proteolysis. **E,** TEM images from WT, *SLP2* KO and *MIC26* KO cells. *SLP2* KO shows accumulation of swollen cristae, while *MIC26* KO shows interconnected cristae arranged in a honeycomb manner. **F,** Cristae number and CJs per mitochondrial section quantified from TEM images. Statistical analysis was performed with one-way ANOVA. *P-value ≤ 0.05, **P- value ≤ 0.01, ***P-value ≤ 0.001.

SLP2 interacts with mitochondrial proteases like YME1L, OMA1 and PARL and forms a large hub of proteases within the IM known as SPY complex (Wai, Saita et al., 2016). We also confirmed the interaction between SLP2 and YME1L (Fig 1C). However, previous study did not identify any YME1L-specific substrate which is regulated by SLP2. Thus, we asked whether the loss of MIC26 in *SLP2* KO cells occurs due to proteolysis via YME1L. To test this, we depleted YME1L in *SLP2* KO cells using shRNA, and indeed found a rescue of the levels of MIC26 in *SLP2* KO cells (Fig 3C). Overall, we found that SLP2 specifically protects MIC26 from proteolysis via YME1L in the MICOS complex (Fig 3D). We conclude that MIC26 is a novel substrate of YME1L which is regulated by SLP2 and well in line with the regulation of proteases by scaffolding proteins. Thus, we identified novel quality control axis of SLP2- YME1L which specifically regulates the steady state levels of MIC26 in MICOS assembly.

### Loss of SLP2 impairs formation of cristae and crista junctions

To determine how SLP2 interaction with MICOS subunits and reduction of MIC26 in *SLP2* KO cells would influence cristae morphology, transmission electron microscopy (TEM) images were acquired from *SLP2* KO cells. *SLP2* KO cells showed swollen cristae and significant reduction in the number of cristae and CJs compared to the control cells (Fig 3E, F). To determine whether cristae defects in *SLP2* KO occur due to reduced MIC26, we tried to express MIC26 in *SLP2* KO cells. However even after multiple trials it was not possible to obtain *SLP2* KO cells with stable MIC26 overexpression, perhaps due to the fact that MIC26 could not to be stabilized without SLP2 until YME1L is present. Therefore, we compared the cristae morphology of *SLP2* KO with *MIC26* KO cells to determine if they show similar cristae defects. *MIC26* KO cells showed branching of cristae which appear like a honeycomb, while compared to wild-type cells the number of cristae were slightly reduced, yet the number of CJs were comparable (Fig 3F). Overall, the cristae defect and the reduction in cristae number in *SLP2* KO was severe compared to *MIC26* KO. This indicates that cristae defects in *SLP2* KO cells do not only arise from loss of MIC26 but rather suggests that SLP2 has an additional role in MICOS assembly and cristae morphology apart from maintaining the stability of MIC26. Overall, we demonstrate that SLP2 is new auxiliary subunit of MICOS complex that could modulate cristae and CJ.

### SLP2 and the MIC13 synergistically regulate MICOS assembly

To determine whether there is any synergistic role of MIC13 and SLP2 in MICOS assembly and cristae morphology, we deleted *SLP2* in *MIC13* KO to generate a double KO (DKO) of *MIC13* and *SLP2*. Steady state levels of MICOS subunits were similar in single *MIC13* KO and *MIC13-SLP2* DKO as the effect was due to MIC13 deficiency. However, MIC26 levels were even more reduced in *MIC13-SLP2* DKO as compared to single *SLP2* KO or *MIC13* KO cells (Fig 4A). The MICOS complex could not assemble fully and ran at a lower size in *MIC13* KO cells due to loss of the MIC10-subcomplex as observed using MIC19 and MIC60 antibodies (Fig 4B). Strikingly, the assembly of MIC19 and MIC60 into the MICOS complex was drastically reduced in *MIC13-SLP2* DKO cells compared to *MIC13* KO. *MIC13-SLP2* DKO cells showed a synergetic outcome on MICOS assembly with the reduced size of MICOS complex as seen in *MIC13* KO cells as well as reduced incorporation of MIC60 and MIC19 into the MICOS complex (Fig 4B). In sum, although single knockout cells lacking *SLP2* or *MIC13* were able to manage MIC60-subcomplex assembly, the double deletion of SLP2 and MIC13 is detrimental to MIC60-subcomplex assembly, showing a synergistic role of SLP2 and MIC13 in mediating the assembly of the MIC60-subcomplex (Fig 4C). In order to determine the influence of such a synergistic regulation of SLP2 and MIC13 on mitochondrial ultrastructure, we analysed the cristae morphology in single and double KOs using TEM. *MIC13* KO displayed onion-like cristae while *SLP2* KO displayed swollen cristae (Fig 4D), both having significantly reduced number of cristae and CJs (Fig 4D, E). *MIC13-SLP2* DKO also had a similar severe phenotype compared to *MIC13* KO cells with significantly reduced numbers of cristae and CJs. Hence, we conclude that synergy between SLP2 and MIC13 is required for the assembly of MIC60- subcomplex and formation of cristae and crista junction.

**Figure 4.**
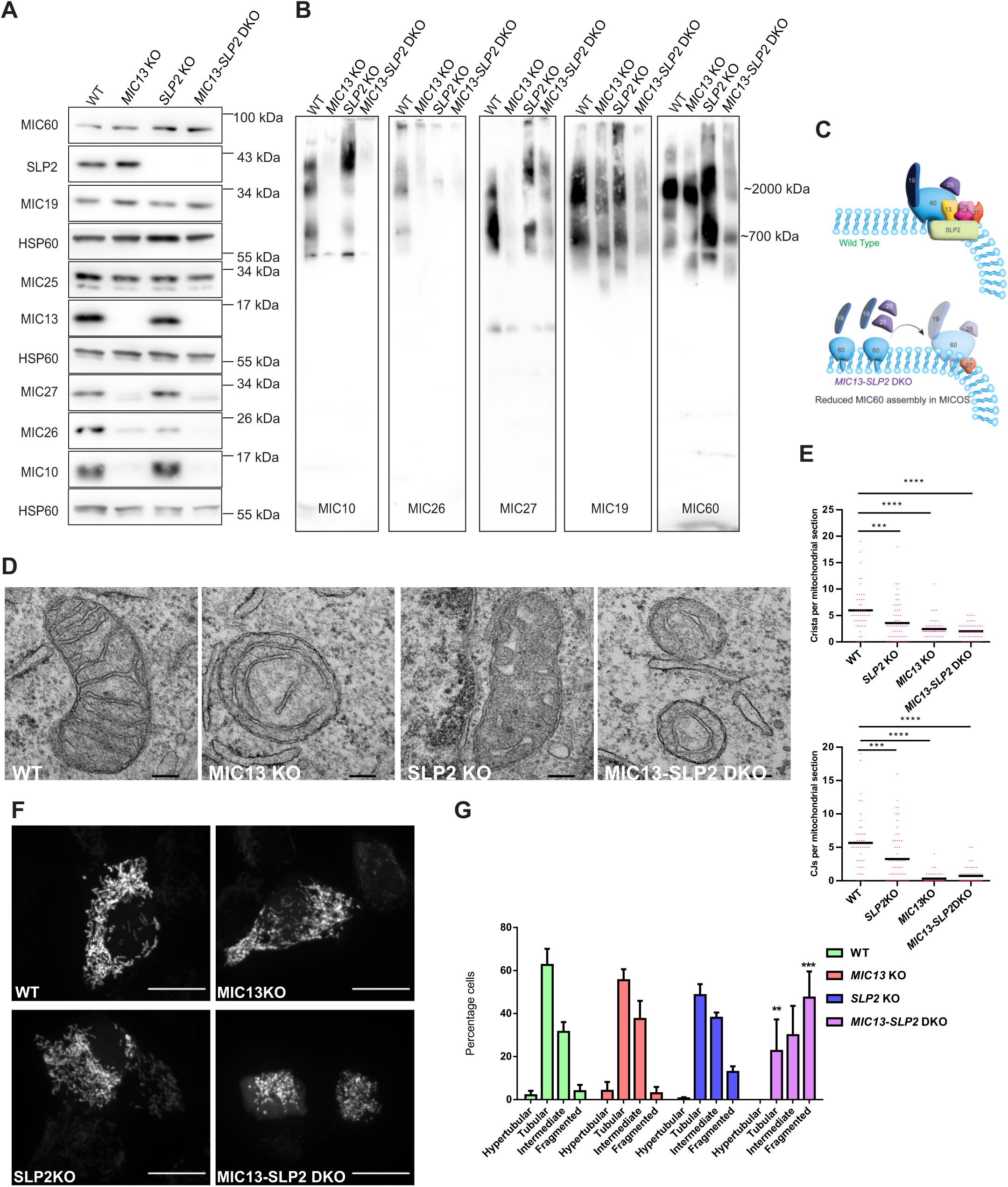
SLP2 and MIC13 synergistically regulate assembly of MIC60-subcomplex. **A**, Assessment of steady state levels of MICOS proteins with western blot from WT, *MIC13* KO, *SLP2* KO and *MIC13-SLP2* DKO cells. **B,** BN-PAGE of isolated mitochondria from WT, *MIC13* KO, *SLP2* KO and *MIC13-SLP2* DKO cells to assess MICOS assembly. *MIC13-SLP2* DKO showed reduced MIC60 assembly in MICOS complex compared to any single KO. **C,** A model depicting that MIC60-subcomplex assembly is dependent on SLP2-MIC13 axis. **D**, TEM images displaying mitochondrial morphology from WT, *MIC13* KO, *SLP2* KO and *MIC13-SLP2* DKO cells. **E,** Quantification of number of cristae and CJs per mitochondrial section obtained from TEM. *P-value ≤ 0.05, **P-value ≤ 0.01, ***P-value ≤ 0.001.

*MIC13-SLP2* DKO showed higher extent of mitochondrial fragmentation compared to either single knockout cells (Fig 4F, G). This could be correlated with lower assembly of MIC60- subcomplex as loss of MIC19 and MIC60 were previously shown to cause mitochondrial fragmentation. SLP2 plays a role in stress-induced mitochondrial hyperfusion (SIMH) (Tondera et al., 2009). To determine whether SLP2-MICOS interaction can influence SLP2-mediated SIMH, mitochondrial morphology upon stress was analysed in single KOs and *MIC13-SLP2* DKO. SIMH was induced by inhibition of protein synthesis using cycloheximide treatment, which showed accumulation of hypertubular mitochondria in WT cells within 2 hours of treatment (Supplementary Fig 2A, B). As expected, *SLP2* KO failed to show any response upon SIMH. *MIC13* KO cells showed hyperfusion similar to WT cells, while *MIC13-SLP2* DKO showed the response which was similar to single *SLP2* KO (Supplementary Fig 2A, B), implying that SLP2-mediated SIMH occurs independent to its interaction with MICOS.

To understand whether SLP2 contributes towards the function of MIC13 to interact with the MIC60-subcomplex and the MIC10-subcomplex, we checked for the interaction of MIC13 with MICOS subunits in the presence and absence of SLP2 by expressing MIC13-FLAG in *MIC13* KO or *MIC13-SLP2* DKO, respectively. MIC13 was able to interact with all MICOS subunits even upon loss of SLP2, although the interaction with MIC26 was reduced due to MIC26 degradation in *SLP2* KO. This suggests that the MIC13-MICOS subunit interactions are independent of SLP2. Additionally, we found a novel interaction of MIC13 with YME1L which was also independent of SLP2 pointing towards a novel SLP2-independent MIC13-YME1L axis in the IM (Supplementary Fig 3). In summary, we found that SLP2 and MIC13 regulate the assembly of MIC60-subcomplex, which was independent to SLP2-mediated SIMH.

### SLP2 selectively regulates the assembly kinetics of MIC60-subcomplex

To mechanistically dissect the role of SLP2 in regulating of MIC60 assembly, we decided to apply the Tet-On system to reintroduce *MIC13* in a time-dependent manner and analyse re-assembly of the MICOS complex over short time scales in *MIC13* KO and *MIC13-SLP2* DKO cells. For this, we generated pLIX403-MIC13-FLAG and stably expressed it in the *MIC13* KO and *MIC13-SLP2* DKO using the lentiviral transduction method. The addition of doxycycline induces the expression of MIC13-FLAG in a time-dependent manner. After eight hours of doxycycline induction, MIC13-FLAG started to express in both *MIC13* KO and *MIC13-SLP2* DKO (Fig 5A). MIC13-FLAG was gradually expressed and incorporated into the MICOS complex of both *MIC13* KO and *MIC13-SLP2* DKO cells as seen by BN-PAGE at different time points after induction of protein expression (Fig 5B). Both MIC10 and MIC27 also progressively assembled in the MICOS complex over time (Fig 5C). The appearance of the MIC10- subcomplex as determined by probing the blots for MIC10 or MIC27 was more rapid and efficient in *MIC13-SLP2* DKO cells compared to the *MIC13* KO cells. This could be due to slightly higher expression of MIC13-FLAG in *MIC13-SLP2* DKO compared to the *MIC13* KO. On the contrary, the incorporation of MIC60 into the MICOS complex was slower in *MIC13- SLP2* DKO cells compared to the *MIC13* KO after induction of MIC13-FLAG. The arrows in the Fig 4C show that MICOS size was recovering yet the amount of MIC60 in MICOS complex took more time to recover (16 hours) in *MIC13-SLP2* DKO (Supplementary Fig 4A, B). Despite the presence of the already assembled MIC10-subcomplex, it took more time for MIC60 to assemble into MICOS complex when SLP2 was missing. The assembly kinetics of the MIC60- subcomplex rather than the MIC10-subcomplex is dependent on SLP2 (Fig 5D), concluding that SLP2 specifically regulates the incorporation of MIC60 into the MICOS complex.

**Figure 5.**
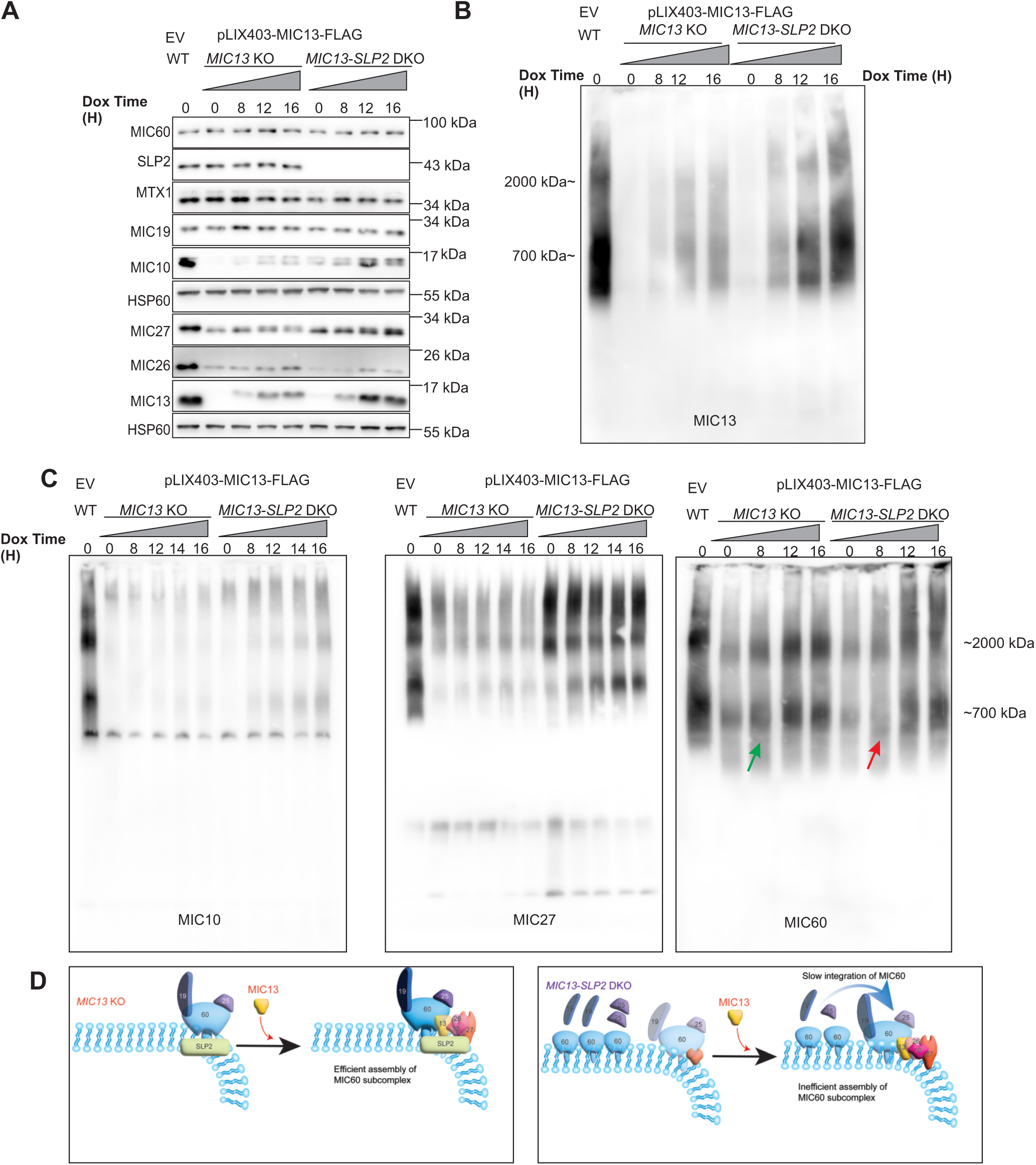
SLP2 specifically regulates assembly kinetics of MIC60. **A**, WT cells stably expressing pLIX403 EV and *MIC13* KO, *MIC13-SLP2* DKO cells stably expressing pLIX403- MIC13-FLAG were treated with 1 μg/ml of doxycycline (Dox) for indicated time points and western blot analysis depicting steady state levels of MICOS proteins upon induction of MIC13- FLAG are shown. **B,** BN-PAGE with isolated mitochondria from WT cells stably expressing pLIX403 EV, and *MIC13* KO and *MIC13-SLP2 DKO* cells stably expressing pLIX403-MIC13-FLAG treated with 1 μg/ml of doxycycline (Dox) for indicated time points showing stable incorporation of MIC13-FLAG in MICOS complex. **C,** Blue native PAGE with isolated mitochondria from WT cells stably expressing pLIX403 EV, and *MIC13* KO and *MIC13-SLP2 DKO* cells stably expressing pLIX403-MIC13-FLAG treated with 1 μg/ml of Dox for indicated time points was probed for MIC10, MIC27 and MIC60 antibody. It shows that kinetics of MIC60 assembly was dependent on SLP2. **D,** A model depicting the assembly kinetics of MIC60 in MICOS complex depends on SLP2.

### MIC13 stabilizes MIC10-subcomplex by regulation of YME1L-mediated proteolysis

Next, we wanted to decipher the specific role of MIC13 in MIC13-SLP2 alliance for assembly of MIC60-subcomplex. However, it has been difficult to identify the exact molecular role of MIC13 because *MIC13* KO was always associated with loss of the MIC10-subcomplex making it hard to differentiate whether the effects were arising due to MIC10 or MIC13 loss. Based on two observations: a novel interaction between YME1L and MIC13 which is independent of SLP2 (Supplementary Fig 3) and an increase in MIC10 levels upon the YME1L downregulation in all the cell lines (Fig 3C), we hypothesized perhaps MIC10 degradation in *MIC13* KO could be YME1L-mediated. We generated cell lines with stable expression of shRNA against YME1L in WT, *SLP2* KO, *MIC13* KO and *MIC13-SLP2* DKO cell lines. Upon knockdown of YME1L, the levels of MIC10 and MIC27 were not only enhanced in *MIC13* KO but also in *SLP2* KO and *MIC13-SLP2* DKO cells showing that both MIC10 and MIC27 are novel YME1L substrates which are regulated independent of SLP2 (Fig 6A). Next to the already described SLP2- dependent YME1L proteolysis of MIC26 (Figure 4), here we identified a second quality control axis of YME1L-mediated proteolysis of MIC10 and MIC27, which is independent to SLP2 but rather dependent on MIC13, demonstrating an intricate mechanism of YME1L-dependent quality control within the MICOS complex. Moreover, YME1L downregulation in *MIC13* KO cells which restored MIC10 levels provides us an exclusive scenario to determine the specific roles of MIC13 that are independent to MIC10. In *MIC13* KO cells, the MIC60-subcomplex shows a lower molecular weight compared to the WT due to loss of the MIC10-subcomplex (Fig 6B). We checked whether the restored levels of MIC10 using YME1L shRNA could promote assembly of MIC10 with the MIC60-subcomplex in *MIC13* KO cells. Indeed, we observed a MIC60 size shift in *MIC13* KO cells upon additional YME1L knockdown showing that the MIC10-subcomplex can assemble with the MIC60-subcomplex despite the absence of MIC13 (Fig 6B). This changes our view on MICOS assembly as it implies that MIC13 is not required to bridge the MIC10- and MIC60-subcomplex, rather it plays an indispensable role in stabilizing the MIC10-subcomplex and controls the quality of the MIC10-subcomplex via YME1L-dependent proteolysis (Fig 6B).

**Figure 6.**
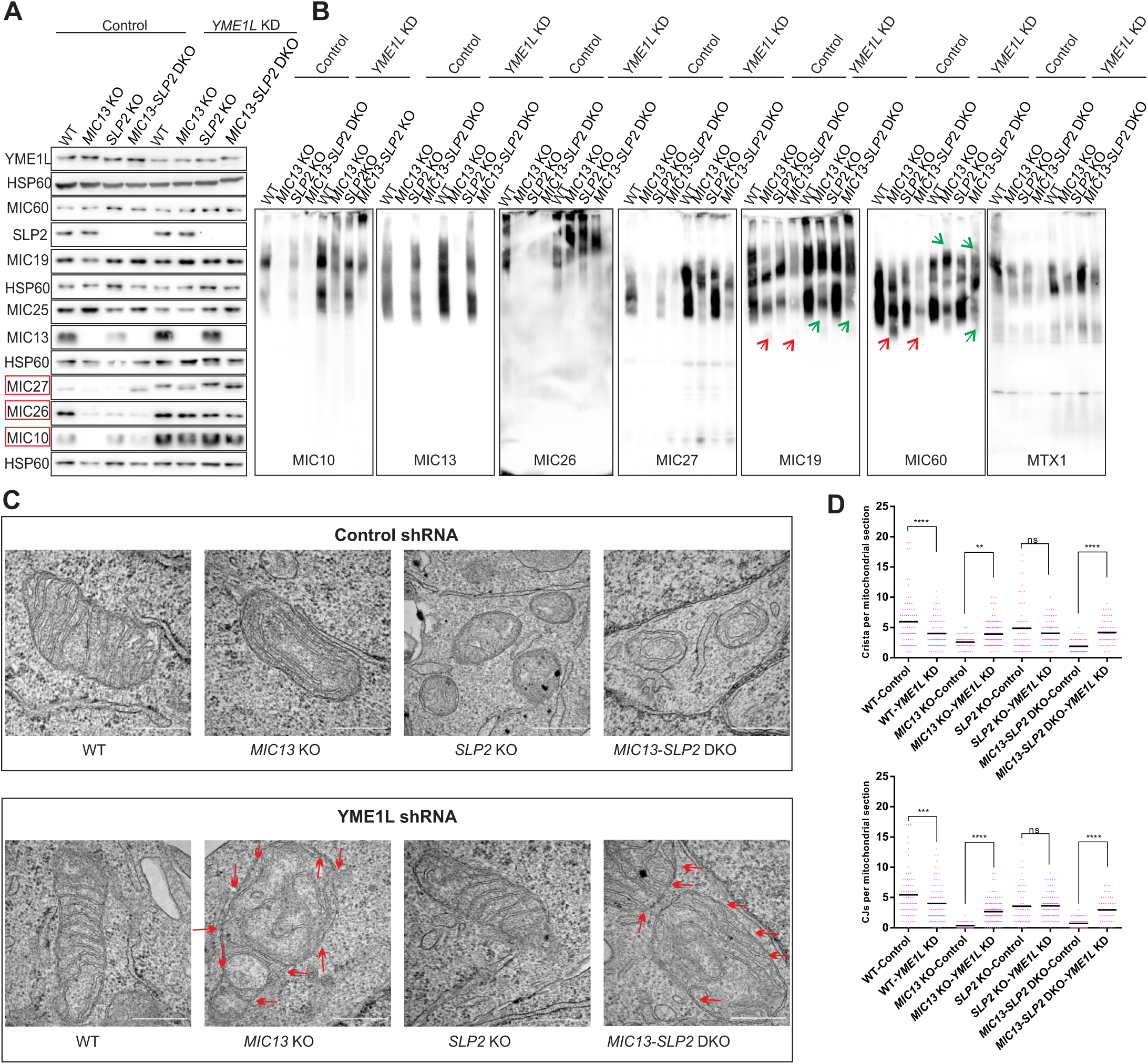
MIC13-YME1L and SLP2-YME1L axes stabilize MIC10- and MIC60-subcomplex assembly in a co-dependent manner. **A**, WT, *MIC13* KO, *SLP2* KO, *MIC13-SLP2* DKO stably expressing pGIPZ-Control shRNA or pGIPZ-YME1L shRNA (knockdown represented as KD) subjected to western blot to assess steady state levels of MICOS proteins. The levels of MIC10, MIC26 and MIC27 were dependent on YME1L-mediated proteolysis. **B,** BN-PAGE with isolated mitochondria from WT, *MIC13* KO, *SLP2* KO, *MIC13-SLP2 DKO* stably expressing pGIPZ-Control shRNA or pGIPZ- YME1L shRNA. Red arrow indicates downshift of MIC60 and green arrow indicates upshift of MIC60 in BN-PAGE. **C,** Mitochondrial cristae morphology accessed using TEM from WT, *MIC13* KO, *SLP2* KO, *MIC13-SLP2* DKO stably expressing pGIPZ-Control shRNA or pGIPZ-YME1L shRNA. Scale bar represents 0.5 µm. Red arrows depict CJs in the mitochondrial section showing a partial beneficial effect on cristae morphology upon YME1L depletion. **D,** Quantification of crista and CJs per mitochondrial section. Outliers were removed with Grubbs’ method and statistical significance was analysed by one-way ANOVA. *P-value ≤ 0.05, **P-value ≤ 0.01, ***P-value ≤ 0.001.

### Stabilization of MIC10-subcomplex could partially promote CJ formation in *MIC13* KO

To determine the interdependence between SLP2 and MIC13 for MIC60 assembly, we wanted to check whether restoration of the MIC10-subcomplex in *MIC13-SLP2* DKO cells could improve the MIC60 assembly into MICOS complex. Even though the steady state levels of MIC60-subcomplex were not affected upon YME1L depletion, the assembly and the shift in MIC60-subcomplex was partially restored in the *MIC13-SLP2* DKO background (Fig 6B). This indicates that the assembled MIC10-subcomplex, in absence of SLP2, can partially provide a ‘docking platform’ for MIC60 and MIC19 to incorporate in the MICOS complex. The partial restoration of MIC60-MIC19 assembly also reflected in the assembly of MIB complex protein MTX1 (Fig 6B). MTX1 is a MIB subunit, together with SAMM50, it forms contact sites between the IM and OM (Huynen et al., 2016). *MIC13-SLP2* DKO cells displayed a drastic reduction in MTX1 assembly, which was restored upon depletion of YME1L (Fig 6B). Together, this data points towards a larger role of SLP2-MIC13 axis in efficient assembly of the MIB complex in association with MIC60-subcomplex.

To further validate the importance of the MIC10-subcomplex promoting the assembly of MIC60 into a functional MICOS and MIB complex, we investigated mitochondrial ultrastructure upon depletion of YME1L in the single and DKO cell lines. Depletion of YME1L from *MIC13* KO and *MIC13-SLP2* DKO cell lines led to a moderate beneficial effect on mitochondrial ultrastructure, as the number of CJs were restored (Fig 6C, D). The images of a single mitochondrion showed some cristae displaying intact CJ where it appeared that onion-shaped cristae unfurled to form nascent CJ. Thus, indicating that stabilized MIC10-subcomplex can promote the formation of nascent CJ in *MIC13* KO or *MIC13-SLP2* DKO (Fig 6C, D).

### SLP2 and MIC13 regulate the formation of MIC60 puncta in IM

We show that MIC60 assembly into the MICOS complex was reduced in *MIC13-SLP2* DKO cells, while the steady state levels of MIC60 was comparable to controls. Therefore, we wanted to determine the status of MIC60 distribution in the IM in these KO cell lines. Using super-resolution stimulated emission depletion (STED) nanoscopy, the distribution of MIC60 in the IM was marked with MIC60-specific antibodies. MIC60 showed punctate pattern with the rail-like arrangement, which is consistent with previous publications (Jans, Wurm et al., 2013, Kondadi et al., 2020, Stoldt, Stephan et al., 2019). This pattern of MIC60 staining resembles the arrangement of CJs in the mitochondria. MIC60 staining in the *MIC13-SLP2* DKO was remarkably different, compared to the WT cells, with less punctate structures and diffuse staining pattern evenly spread along the IBM. The staining of MIC60 in single *MIC13* KO and *SLP2* KO was also perturbed and the rail-like pattern was less prominent as the MIC60 spots were also equally distributed within the mitochondria (Fig 7A). MIC60 could be present on the inner cristae stacks that are accumulated in *MIC13* KO and *SLP2* KO. Even though the single KOs could manage MIC60 puncta formation, the combined loss of SLP2 and MIC13 was detrimental to MIC60 puncta formation, demonstrating a critical role of SLP2 in dictating formation of MIC60 puncta perhaps by facilitating the formation of a conducive lipid microenvironment. Stomatins are shown to create the microdomains in the lipid-bilayer. Therefore, we propose that SLP2-MIC10-subcomplex acts as ‘seeder’ for the formation of MIC60 puncta in IM and thereby promote formation of CJ.

**Figure 7.**
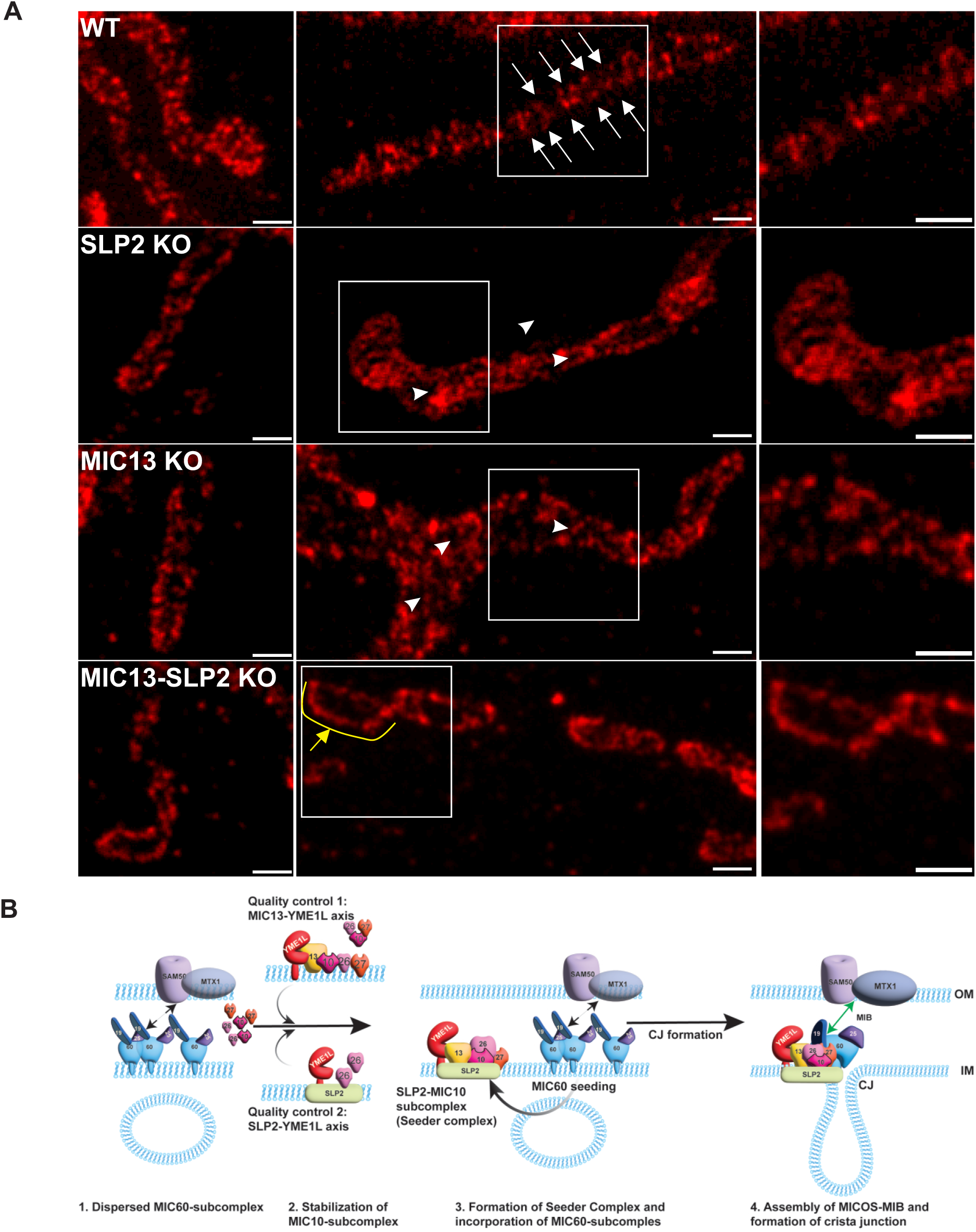
MIC13-SLP2 axis is required for formation of MIC60 punctae. **A**, STED nanoscopy images from WT, *MIC13* KO, *SLP2* KO and *MIC13-SLP2 DKO* cells displaying MIC60 punctae. White arrows indicate individual MIC60 punctae in a rail-like arrangement in WT cells. Arrow heads depict perturbed MIC60 punctae in *MIC13* KO and *SLP2* KO. Yellow arrow with curve depicts dispersed MIC60 punctae in *MIC13-SLP2* DKO cells. **B,** A comprehensive schematic model of CJ formation illustrating the novel quality control process orchestrated by the MIC13-YME1L and SLP2-YME1L axis, which facilitates the formation of the SLP2-MIC10-subcomplex, known as the “seeder complex.” The seeder complex promotes the ‘seeding’ or assembly of the MIC60-subcomplex and the essential MIB protein MTX1, consequently playing a crucial role in defining the formation of MIC60 puncta and the MICOS- MIB complex. This mechanistically promote the formation of CJ and contact between IM and OM.

Overall based on our results, we propose a ‘seeder’ model for the assembly of MICOS complex which shows interdependence of the assembly of MIC60-subcomplex and the MIC10- subcomplex on each other and on the scaffolding protein, SLP2 (Fig 7B). The arrangement of the MIC60 was dispersed upon loss of both SLP2 and MIC13. The preliminary MIC60- subcomplex is dispersed in the IM and unable to contact MIB components. The MIC10- subcomplex is intricately stabilized and assembled by two differentially regulated quality control axes dependent on YME1L. The ‘seeder’ complex is formed by association of SLP2 and MIC10-subcomplex, which provides a docking platform for further assembly of MIC60- subcomplex into holo-MICOS complex and MIB complex. This is shown by formation of MIC60- specific puncta in the presence of both SLP2 and MIC13 and leading to reattachment of the cristae stacks to IM and formation of nascent CJ as well contact site between IM and OM. The concomitant reappearance of CJ upon stabilization of MIC10-subcomplex provides mechanistic insights for formation of nascent CJ.

## Discussion

MIC13 being an integral component of the mitochondrial IM has been vital for cristae morphogenesis and loss of MIC13 leads to severe mitochondrial hepato-encephalopathy. Although previous research has extensively shown the consequence upon MIC13 depletion, molecular mechanisms leading to the phenotypes are largely unknown. Here we determined the interactome of MIC13 to obtain novel insights about the unidentified molecular role of MIC13. The interactome data mostly contained proteins belonging to MICOS-MIB complex or the known interactor of these complexes, implying a crucial role of MIC13 in MICOS regulation. Among few novel interactors, SLP2 was one of the most highly enriched protein. SLP2 belongs to the SPFH (stomatin, prohibitin, flotillin, HfLC/K) superfamily which is composed of scaffolding proteins that can locally define the protein and lipid configuration of a cellular membrane (Wai et al., 2016). Though an interesting possibility, a direct role of SLP2 in MICOS assembly and cristae dynamics was not reported earlier. Using KOs of individual MICOS subunits and stable overexpression cell lines, we determined that SLP2 not only interacts with MIC13 but also with each MICOS subunit and forms a large interaction hub of SLP2-MICOS in the mitochondrial IM. Our results show novel multi-layered roles of SLP2 in regulating the MICOS assembly and CJ formation. SLP2 was identified as a new cristae and CJ modulator as loss of SLP2 leads to defects in cristae and CJ morphology. Based on these results, we propose SLP2 as an auxiliary subunit of MICOS. SLP2 was required for stability of MIC26 and its assembly into MICOS complex. It protects MIC26 from YME1L-mediated proteolysis. SLP2 was earlier shown to form proteolytic hub with mitochondrial proteases PARL, YME1L and perhaps OMA1 known as SPY complex. However, no earlier known substrates of YME1L were shown to be regulated by the SPY complex (Wai et al., 2016). We show here that MIC26 is a novel substrate of YME1L which is specifically regulated by the SLP2-YME1L axis. Recently, we have reported a pathological mutation in *MIC26* that causes lethal mitochondrial disease with progeria-like phenotypes (Peifer-Weiß et al., 2023). This MIC26 variant with loss of the last twenty amino acids at the C-terminus is prone to faster degradation such that the mutant behaves like a loss-of-function variant. Perhaps MIC26 degradation in patient cells is the consequence of enhanced SLP2-YME1L-mediated proteolysis. *MIC26* mutations were also associated with recessive mitochondrial myopathy, lactic acidosis, cognitive impairment and autistic features (Beninca, Zanette et al., 2021b). Further study of SLP2-YME1L axis to regulate MIC26 could provide the patho-mechanisms in MIC26-associated pathologies.

Although SLP2 showed a specific regulation of MIC26, our interaction data and the fact that *SLP2* KO shows more severe cristae defects compared to *MIC26* KO pointed towards possible broader roles of SLP2 in MICOS assembly and cristae architecture. Peculiarly, the DKO of *MIC13* and *SLP2* showed more additive defects compared to each single KO with respect to the MIC60 assembly into the MICOS complex. This indicated that MIC13 and SLP2 function in unison to regulate the incorporation of MIC60 into the MICOS complex. The time-dependent restoration of MIC13 in *MIC13-SLP2* DKO cells showed a considerably slower kinetics of the assembly of MIC60 into MICOS complex compared to *MIC13* KO. This demonstrated a critical role of SLP2 in regulating the kinetics of MIC60-subcomplex assembly and CJ formation.

To dissect the specific role of MIC13 in the regulation of MIC60 assembly in SLP2-MIC13 axis, we wanted to differentiate whether the phenotypes arise from MIC10 or MIC13 as *MIC13* KO is always associated with loss of MIC10-subcomplex. Based on our observations that MIC13 interacts with YME1L and MIC10 levels are enhanced in YME1L downregulation, we checked whether MIC10 levels are regulated by YME1L in *MIC13* KO cells. The downregulation of YME1L in *MIC13* KO could not only restore the steady state levels of MIC10 but also its incorporation into the MICOS complex. Since MIC10 could be integrated in MICOS complex in absence of MIC13, we conclude that while MIC13 in principle is dispensable for bridging MIC10- and MIC60-subcomplexes, it rather plays an important role in stabilizing the MIC10- subcomplex via inhibiting YME1L-specific proteolysis. The presence of nascent CJ upon depletion of YME1L in *MIC13* KO showed that the assembled MIC10-subcomplex could form CJ even in the absence of MIC13. The cristae in this scenario appeared as if the onion-like cristae of *MIC13* KO unfurled and reattached to IM to form nascent CJ, providing a model for CJ formation. MIC10 is known to oligomerize in a wedge-like shape that causes membrane bending and CJ formation (Barbot et al., 2015, Bohnert et al., 2015). However, here question remains whether this CJ restoration is sufficient to restore MIC13-specific mitochondrial defects or the associated pathology. We also observe that MIC27 showed a band at a lower apparent molecular weight in *MIC13* KO. The identity of the lower MIC27 is not yet known but it was interesting to note that conversion of MIC27 to a lower size was MIC13-dependent and was not influenced by YME1L depletion. This is because even when the steady state levels of MIC27 was restored upon downregulation of YME1L in *MIC13* KO, the status of the MIC27 band size was not restored. The YME1L-dependent restoration of MIC10 presents a new tool to distinguish the MIC13-specific functions, which has direct implications in study of the MIC13- associated disease mechanism and in development of future therapeutics.

Using YME1L as a tool, we restored the MIC10 levels in *MIC13-SLP2* DKO and checked for MIC60 incorporation into MICOS complex. Here, we found that restoration of MIC10 could partially restore the incorporation of MIC60 into the MICOS complex in the *MIC13-SLP2* DKOs. This showed an interdependence of SLP2 and MIC10-subcomplex for the MIC60 assembly, where SLP2 and the MIC10-subcomplex that is stabilized by MIC13 act as a ‘seeder’ for incorporation of MIC60 into the holo-MICOS complex. We termed SLP2-MIC10-subcomplex as a ‘seeder’ complex. In agreement, the super-resolution STED nanoscopy showed a dispersed arrangement of MIC60 in the IM in *MIC13-SLP2* DKO compared to a normal rail-like punctate arrangement in the control cells. This implies that MIC60 in the absence of SLP2 and MIC13 could not be confined to MICOS puncta but rather remains dispersed. SLP2 and MIC13 direct the site for MIC60 puncta formation and thereby the CJ formation. Perhaps SLP2 role is to generate special lipid environment for efficient MIC60-subcomplex assembly and its association to its docking partner, MIC10-subcomplex. In conclusion, both SLP2 and MIC13 work together to generate a conducive environment for efficient incorporation of MIC60 into MICOS-MIB complex and thereby CJ formation.

Initially it has been thought that MICOS assembly follows a hierarchy where MIC60 subcomplex assembly happens prior to MIC10-subcomplex and MIC60 act as a master regulator (Ott, Dorsch et al., 2015, Zerbes, Hoss et al., 2016). However, our results show an interdependence between these two steps: wherein kinetics of MICOS subcomplex assembly progresses in an interdependent manner, concurrently SLP2 promotes the efficient assembly of MIC60-subcomplex. Hence, we propose the ‘seeder’ model for the assembly of the MICOS- MIB complex. We identified two differentially regulated quality control processes, SLP2-YME1L and the MIC13-YME1L axis, that determine the stability and assembly of MIC10-subcomplex. Once the MIC10-subcomplex is stabilized, the collaboration between MIC10-subcomplex and the auxiliary MICOS subunit SLP2 (seeder complex) acts as a ‘seeder’ for formation of MIC60- specific puncta and thereby incorporation of MIC60-subcomplex into MICOS-MIB complex. Therefore, the seeder complex dictates the site for formation of MIC60 puncta and hence the morphogenesis of CJ and contact site between the IM and OM.

The function of CJ and contact sites are not fully understood. Defects in the MICOS subunits are shown to affect several important cellular functions including import of mitochondrial protein, biogenesis and transfer of phospholipids, mtDNA organization, apoptosis, mitophagy, mitochondrial transport, mitochondrial translation, mitochondrial morphology and inflammation (Anand et al., 2021). The role of SLP2 and newly identified quality control axes in process of MICOS-MIB could provide new insights into many of these cellular processes in future studies.

The quality control of MICOS assembly is intricately regulated by YME1L-mediated proteolysis at different steps. SLP2-YME1L axis regulates the MIC26 stability and function. A Novel MIC13-YME1L axis reported here, on the other hand, regulates MIC10, MIC26 and MIC27 levels independent of SLP2. MIC60 and SAMM50 were also proteolytic substrates of YME1L during *MIC19* downregulation (Li, Ruan et al., 2016, Tang et al., 2020) showing the complexity of regulation of MICOS assembly. Another IM protease, OMA1 also associates with the MICOS complex (Viana, Levytskyy et al., 2021) and proteolytically regulates MIC19 assembly in the MIB complex (Tang et al., 2020). Overall, our study deciphers a novel quality control mechanism in regulating MICOS and MIB assembly and expands our understanding on the factors regulating cristae morphogenesis.

### Materials and methods Cell culture

Flp-In T-REx HEK293 and HeLa cells were cultured in Dulbecco’s modified Eagle medium (DMEM) with 1g/L glucose and sodium pyruvate (PAN-Biotech, P04-01500) supplemented with 1% stable glutamine (P04-82100), 1% penic illin-streptomycin (Sigma-Aldrich, P4333- 100ml), 10% fetal bovine serum (FBS) (Capricorn Scientific, FBS-11A). Plat-E and HEK293FT cells were cultured in DMEM high glucose medium (PAN-Biotech, P04-03500) supplemented with 10%FBS, 1% stable glutamine, 1% sodium pyruvate (Gibco, 11360070). GIPZ-Control-Flp-In T-REx HEK293 and GIPZ-YME1LshRNA- Flp-In T-REx HEK293 cells were cultured in DMEM 1g/L glucose medium containing sodium pyruvate supplemented with 1% stable glutamine, 10% FBS, 1% penicillin-streptomycin and 1% Non-Essential Amino Acids Solution (NEAA) (PAN-Biotech, P08-32100). All cells were cultured in at 37°C with 5% CO2. All cell line generated are listed in Supplementary Table 2.

### Co-immunoprecipitation coupled mass spectrometry

Co-IP was performed with Protein A Sepharose CL-4B beads (Invitrogen, 101041) and affinity purified MIC13 antibody was linked to the beads. Isolated mitochondria from Flp-In T-REx HEK293 WT or *MIC13* KO were solubilized with isotonic buffer (150 mM NaCl, 10 mM Tris/HCl (pH 7.5), 5 mM EDTA, 1x protease inhibitor cocktail) supplemented with 5 µl of 10% Digitonin (2g/g of protein) and added to the beads with subsequent incubation in 4°C under rotation conditions. Beads were washed several times, transferred into a new tube in 10mM Tris, pH 7.4. Beads were resuspended in 50 µl 6M GdmCl, 50 mM Tris/HCl, pH 8.5 and incubated at 95°C for 5 min. Sample were diluted with 25 mM Tris/HCl, pH 8.5, 10% acetonitrile to obtain a final GdmCl concentration of 0.6 M. Proteins were digested with 1 µg Trypsin (sequencing grade, Promega) overnight at 37°C under gentle agitation. Digestion was stopped by adding trifluoroacetic acid to a final concentration of 0.5%. Peptides were loaded on multi-stop-and-go tip (StageTip) containing six C18 discs. Purification and elution of peptides was performed as described in Kulak, *et al* (Kulak, Pichler et al., 2014). Peptides were eluted in wells of microtiter plates and peptides were dried and resolved in 1% acetonitrile, 0.1 % formic acid. Liquid chromatography/ mass spectrometry (LC/MS) was performed on Thermo Scientific™ Q Exactive Plus equipped with an ultra-high performance liquid chromatography unit (Thermo Scientific Dionex Ultimate 3000) and a Nanospray Flex Ion-Source (Thermo Scientific). Peptides were loaded on a C18 reversed-phase precolumn (Thermo Scientific) followed by separation on a with 2.4 µm Reprosil C18 resin (Dr. Maisch GmbH) in-house packed picotip emitter tip (diameter 100 µm, 30 cm long from New Objectives) using an gradient from mobile phase A (4% acetonitrile, 0.1% formic acid) to 60 % mobile phase B (99% acetonitrile, 0.1% formic acid) for 90 min with a flow rate 350 nl/min. MS data were recorded by data dependent acquisition Top10 method selecting the most abundant precursor ions in positive mode for HCD fragmentation. Lock mass option (Olsen, Godoy et al., 2005) was enabled to ensure high mass accuracy between multiple runs. The Full MS scan range was 300 to 2000 m/z with resolution of 70000, and an automatic gain control (AGC) value of 3*106 total ion counts with a maxim al ion injection time of 160 ms. Only higher charged ions (2+) were selected for MS/MS scans with a resolution of 17500, an isolation window of 2 m/z and an automatic gain control value set to 105 ions with a maximal ion injection time of 150 ms. Selected ions were excluded in a time frame of 30s following fragmentation event. Fullscan data were acquired in profile and fragments in centroid mode by Xcalibur software. For data analysis MaxQuant 1.6.1.0 (Cox and Mann, 2008, Nat. Biotechnology), Perseus 1.5.6.0 (Tyranova et al 2016) and Excel (Microsoft Office 2013) were used. N-terminal acetylation (+42.01) and oxidation of methionine (+15.99) were selected as variable modifications and carbamidomethylation (+57.02) on cysteines as a fixed modification. The human reference proteome set (Uniprot, July 2017, 701567 entries) was used to identify peptides and proteins with a false discovery rate (FDR) less than 1%. Minimal ratio count for label-free quantification (LFQ) was 1. Reverse identifications and common contaminants were removed and the data-set was reduced to proteins that were identified in at least 5 of 7 samples in one experimental group. Missing LFQ or IBAQ values were replaced by random background values. Significant interacting proteins were determined by permutation-based false discovery rate (FDR) calculation and students T-Test. The mass spectrometry proteomics data have been deposited to the ProteomeXchange Consortium via the PRIDE [1] partner repository with the dataset identifier PXD044968.

### Proximity ligation assay

PLA was carried out with Duolink® In Situ Red Starter Kit Mouse/Rabbit (Sigma-Aldrich, DUO92101-1KT) following manufacturer’s protocol with minor modifications. Briefly, HeLa cells were fixed using 4% paraformaldehyde (Sigma-Aldrich, P6148) for 20 mins in room temperature and washed with PBS 3x with subsequent permeabilization with 0.15% Triton X- 100 (Sigma-Aldrich, T8787-100ML) in room temperature for 15 mins followed by PBS wash. Permeabilized cells were blocked with blocking solution (provided in kit) for 1 hour in 37°C. Blocking solution was removed and primary antibodies with 1:100 dilution ratio was added to the samples and incubation was carried out at 37°C for 2 hours. Following primary antibodies were used: MIC10 (Abcam, 84969), MIC13 (custom made by Pineda (Berlin) against human MIC13 peptide CKAREYSKEGWEYVKARTK), MIC19 (Proteintech, 25625-1-AP), MIC25 (Proteintech, 20639-1-AP), MIC26 (Thermofisher Scientific, MA5-15493), MIC27 (Sigma-Aldrich, HPA000612-100UL), MIC60 (Abcam, ab110329), SLP2 (Abcam, ab102051), SLP2 (OriGene, TA808240), Mt-CO2 (Abcam, ab110258). Subsequent ligation of PLA probes and amplification of circular DNA probes was carried out following manufacturer’s protocol. PLA signals were visualized in PerkinElmer spinning disc confocal microscope equipped with a 60× oil objective.

### CRISPR-Cas9 knockout generation

CRISPR-Cas9 double nickase plasmid (Santa Cruz Biotechnology, SLP2: sc-403638-NIC, MIC26: sc-413137-NIC, MIC60: sc-403617-NIC, MIC25: sc-413621-NIC, MIC19: sc-408682-NIC, MIC10: sc-417564-NIC, MIC27: sc-414464-NIC) was transfected with GeneJuice (Sigma-Aldrich, 70967-3) in Flp-In T-REx HEK293 WT, *MIC13* KO parental lines to generate KO and double KO cell lines. Briefly, cell lines were transfected at 60-70% confluency with 1 µg of double nickase plasmid and incubated for 48 hours followed by 2.5 µg/ml puromycin selection for 24 hours with subsequent single cell sorting based on green florescent protein (GFP) expression using flow cytometry in 96 well plate. Cells were incubated and upon visible colonies, cells were sub-cultured and KO screen was performed with western blotting. Cell lines showing no immune reactivity to respective antibodies were termed as KOs or double KOs.

### Molecular cloning

Human SLP2-HA ORF (Sino Biologicals, HG16147-CY) was cloned into pMSCVpuro vector using Gibson assembly cloning kit (NEB, E2611L), following manufacturer’s protocol. HA tag was replaced with MYC tag with Site Directed and Ligation Independent Mutagenesis (SLIM) (Chiu, Tillett et al., 2008) to generate pMSCV-SLP2-MYC. Human MIC13-Flag from pMSCV puro MIC13-FLAG (Urbach et al., 2021) was cloned into pLIX403 (Addgene, 41395) with Gibson assembly following manufacturers protocol. Primer sequences for Gibson assembly and SLIM are provided in Supplementary Table 3.

### Generation of stable cell lines

For retroviral transduction, Plat-E cells were transfected with 1 µg of pMSCV-MIC13Flag or pMSCV-SLP2MYC and 1 µg of pVSV-G with 3.5 µL of GeneJuice per wells of 6-well plate. After 72 hours incubation, viral supernatant was added to the target cells. Media was replaced with puromycin containing media (2.5 µg/ml) 48 hours of transduction. Puromycin selection was carried out for 2 weeks and successful expression of exogenous protein was validated with western blot. For lentiviral transduction, HEK293FT cells were transfected with 1ug of pLIX403 EV or pLIX403-MIC13-Flag or pGIPZ-non-silencing-Control-shRNA (Horizon Discovery, RHS4346) or pGIPZ-YME1L-shRNA (Horizon Discovery, RHS4430-200157017, RHS4430-200215861, RHS4430-200221198, RHS4430-200268420, RHS4430-200273633, RHS4430-200280144) along with 1µg of psPAX2 (Addgene, 12260) and pMD2.G (Addgene, 12259) was transfected using GeneJuice. 72 hours post transfection, viral supernatant was collected and added on target cell lines. Media was replaced with puromycin (2.5 µg/ml) containing media and selection was carried out for about 2 weeks. Successful exogenous protein expression or knockdown was confirmed with western blotting.

### SDS PAGE and Western blot

Cells were grown in 6-well dishes and harvested with cold PBS upon 70-90% confluency followed by protein extraction by RIPA lysis. Protein concentration was determined by Lowry method (Bio-Rad, 5000113, 5000114, 5000115) and samples were prepared using Laemmli loading buffer. Proteins were separated using 10% or 15% SDS-PAGE with subsequent transfer on nitrocellulose membrane (Amersham, 10600004) followed by 1 hour of blocking using 5% skimmed milk (Carlroth, 68514-61-4). Membranes were incubated overnight in 4°C under shaking conditions in primary antibodies: MIC10 (Abcam, 84969), MIC13 (custom made by Pineda (Berlin) against human MIC13 peptide CKAREYSKEGWEYVKARTK), MIC19 (Proteintech, 25625-1-AP), MIC25 (Proteintech, 20639-1-AP), MIC26 (Thermofisher Scientific, MA5-15493), MIC27 (Sigma-Aldrich, HPA000612-100UL), MIC60 (Abcam, ab110329), SLP2 (Abcam, ab102051), beta-tubulin (Cell Signalling Technology, 2128S), HSP60 (sigma, SAB4501464), Mt-CO2 (Abcam, ab110258), YME1L (Proteintech, 11510-1-AP), MTX1 (Abcam, ab233205). Following primary antibody incubation, membranes were washed in TBST and probed with Goat anti-mouse IgG HRP-conjugated antibody (Abcam, ab97023) or goat anti-rabbit IgG HRP-conjugated antibody (Dianova, 111-035-144). Chemiluminescent signal was recorded with VILBER LOURMAT Fusion SL (Peqlab) and quantification was performed with ImageJ.

### Mitochondria Isolation

Cells were grown in 15 cm dishes and scrapped in cold PBS and pelleted at 500g for 5 mins. Cell pellets were resuspended in isotonic buffer (220 mM mannitol, 70 mM sucrose, 1 mM EDTA, 20 mM HEPES (pH 7.5) and 1 × protease inhibitor cocktail (Sigma-Aldrich, 05056489001)) with 0.1% bovine serum albumin (BSA) (Pan-Biotech, P06-1394100). Cells were mechanically homogenized using syringe with 26G cannula for 15 strokes. Cell homogenate was centrifuged at 1000g for 10 mins, supernatant was collected in fresh tube and further centrifuged at 10,000g for 10 mins at 4°C to obtain crude mitochondrial fractions. Crude mitochondrial pellets were resuspended in isotonic buffer and Lowry assay was performed to determine the concentration. Crude mitochondrial fractions were aliquoted, centrifuged at 10,000g for 5 mins and pellets were resuspended in freezing buffer (300 M trehalose, 10 mM KCl, 1 mM EDTA, 10 mM HEPES and 0.1% BSA) and stored in -80°C until further processing.

### Co-immunoprecipitation

Mitochondrial aliquots of 500 µg were pelleted by centrifugation and re-suspended in isotonic buffer (150 mM NaCl, 10 mM Tris/HCl (pH 7.5), 5 mM EDTA, 1x protease inhibitor cocktail) with 10 µl of 10% Digitonin (2g/g of protein) and solubilized for 10 mins on ice. Solubilized proteins were centrifuged at 21,000g for 20 mins and 10% of the supernatant was separated as input fraction. Remaining supernatant were incubated with anti-Flag M2 affinity beads (Sigma) or MYC-Trap agarose beads (ChromTech) overnight in 4°C under rotation. Beads were centrifuged at 3700g for 1 min at 4°C and 300 µl supernatant was stored as unbound fraction. Beads were further washed (4x) with isotonic buffer with 0.01% digitonin. Proteins were eluted with Laemmli buffer without beta-mercaptoethanol at 65°C for 10 min with subsequent addition of 1 µl beta-mercaptoethanol and subjected to SDS-PAGE with subsequent western blotting.

### Visualization of native protein complexes with blue native PAGE

Mitochondrial aliquots of 150ug were centrifuged and pellets were resuspended in 15 µl of solubilization buffer (50 mM NaCl, 2 mM aminohexanoic acid, 50 mM imidazole/HCl pH 7, 1 mM EDTA, protease inhibitor cocktail) with 3 µl of 10% digitonin (2g/g of protein) and incubated on ice for 10 mins. Samples were centrifuged at 21,000g for 10 mins at 4°C and supernatant was collected in fresh tube followed by addition of 50% glycerol and 1.5 μL of 1% Coomassie brilliant blue G-250. Samples were loaded in 3-13% gradient gel and subsequently transferred on methanol activated PVDF membrane. Membranes were blocked in 5% skimmed milk for 1 hour and incubation was carried out overnight in 4°C under shaking conditions with primary antibodies: MIC10 (Abcam, 84969), MIC13 (custom made by Pineda (Berlin) against human MIC13 peptide CKAREYSKEGWEYVKARTK), MIC19 (Proteintech, 25625-1-AP), MIC25 (Proteintech, 20639-1-AP), MIC26 (Thermofisher Scientific, MIC27 (Sigma-Aldrich, HPA000612-100UL), MIC60 (Abcam, ab110329), SLP2 (Abcam, ab102051), MTX1 (Abcam, ab233205). Primary antibodies were washed 3x with TBST and incubated with Goat anti-mouse IgG HRP-conjugated antibody (Abcam, ab97023) or goat anti-rabbit IgG HRP-conjugated antibody (Dianova, 111-035-144) diluted to 1:10000 in 5% skimmed milk in TBST. Chemiluminescent signal was recorded with VILBER LOURMAT Fusion SL (Peqlab) and quantification was performed with ImageJ.

### Mitochondria morphology analysis

Flp-In T-REx HEK293 cells were transfected with 1ug of mitochondrially targeted GFP (Mito-GFP) with along with 3.5 µL of GeneJuice. 24 hours post transfection, cells were treated with 10 µM cycloheximide and incubated for 2 hours at 37°C in CO2 incubator. Media was removed following PBS washing three times. Cells were fixed using 4% paraformaldehyde (Sigma-Aldrich, P6148) for 20 mins in room temperature and washed with PBS 3 times. GFP signals were visualized in PerkinElmer spinning disc confocal microscope equipped with a 60× oil objective. Cells were classified as hypertubular, tubular, intermediate or fragmented based on the majority of mitochondrial population present in the particular cell. Cells classified as hypertubular contained large interconnected tubular mitochondrial networks. Cells classified as intermediate contained a comparable ratio of short tubes or fragmented mitochondria, while cells classified as tubular and fragmented contained mostly long tubular and very short mitochondria fragments, respectively.

### Stimulated emission depletion (STED) super-resolution nanoscopy

Cells were fixed and permeabilized as described earlier (PLA assay). Permeabilized cells were blocked with 5% goat serum and primary antibody incubation was carried out with 1:100 rabbit anti-MIC60 antibody (custom-made, Pineda (Berlin)), which was generated using the peptide CTDHPEIGEGKPTPALSEEAS against human MIC60, overnight at 4°C and 1:100 Aberrior STAR 635P goat-anti-rabbit (2-0012-007-2) secondary antibody incubated at room temperature for 1 h. STED imaging was performed with the Leica SP8 laser scanning confocal microscope coupled with a STED module. Initially, imaging of 80-nm gold particles (BBI Solutions) was carried out in reflection mode for correct alignment of excitation and depletion laser. A 93x glycerol (N.A = 1.3) objective was used with the pinhole set to 0.6 Airy units and a white light laser excitation wavelength of 633 nm was used for sample excitation. STED depletion was carried out with a pulsed STED depletion laser beam of 775 nm wavelength. A hybrid detector (HyD) was used for signal detection in the range from 643 to 699 nm. 13x magnification was used to acquire images covering a field of view of 9.62 x 9.62 µm. No image processing was performed except smoothing carried out with Fiji software.

### Electron microscopy

Cells were cultured in petri dishes until about 80% confluency was reached and chemical fixation was carried out using 3% glutaraldehyde buffered with 0.1 M sodium cacodylate, pH 7.2, followed by cell scrapping and centrifugation. Cell pellets were washed with 0.1 M sodium cacodylate and embedded in 2% agarose. Cell staining was performed with 1% osmium tetroxide for 50 mins with subsequent incubation in 1% uranyl acetate/1% phosphotungstic acid for 1 hour. Samples were further dehydrated with graded acetone series and embedded in spur epoxy resin for polymerization at 65°C for 24 hours. Ultrathin sections of samples were prepared with microtome and images were captured with transmission electron microscope (Hitachi, H7100) at 75V equipped with Bioscan model 792 camera (Gatan) and analyzed with ImageJ software. The images were randomized and the data was analyzed in a double-blind manner by two scientists. Data analysis was carried out by GraphPad prism. Statistical analysis includes one-way Annova test, outlier test was performed with Grubb’s test where indicated using GraphPad Prism.

## AUTHOR CONTRIBUTIONS

Ritam Naha: Investigation, methodology, data curation, formal analysis, visualization, writing— original draft, writing—review and editing.

Rebecca Strohm: Investigation, methodology, data curation, formal analysis. Jennifer Urbach: Investigation, methodology, formal analysis.

Ilka Wittig: investigation, methodology, formal analysis, data curation.

Arun Kumar Kondadi: formal analysis, supervision, validation, funding acquisition, writing— review and editing.

Andreas S. Reichert: supervision, funding acquisition, writing—review and editing.

Ruchika Anand: conceptualization, data curation, formal analysis, supervision, methodology, funding acquisition, project administration, writing—original draft, writing—review and editing.

## Acknowledgements

Authors would like to thank Andrea Borchardt, Tanja Portugall for their technical assistance in cloning and electron microscopy. We further thank Anny Garces Palacio as student assistant for performing microscopy experiments. Electron microscopy was performed at the Core facility for electron microscopy (CFEM) at the medical faculty of the Heinrich Heine University Düsseldorf. The STED imaging experiments were performed at the Centre for Advanced Imaging (CAi) at Heinrich Heine University Düsseldorf. The research was supported by funding from Medical faculty of Heinrich Heine University Düsseldorf, Foko-02/2015 (RA & ASR), FoKo-2020-71 (RA), FoKo 2022-11 (AKK) and Deutsche Forschungsgemeinschaft (DFG) grant, AN 1440/3-1 (RA), AN 1440/4-1 (RA), KO 6519/1-1 (AKK) and SFB 1208 project B12 (ID 267205415) to ASR.

