## Supplementary figures and images for "MIC13 and SLP2 seed the assembly of MIC60-subcomplex to facilitate crista junction formation"

### Supplementary Fig 1

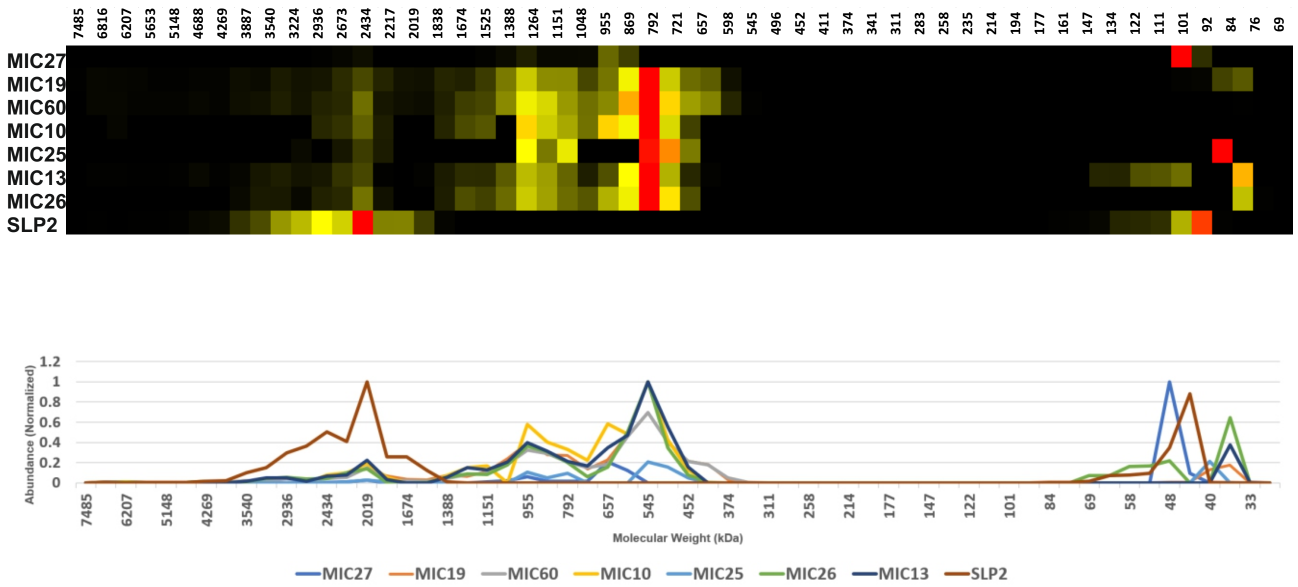
