## Supplementary Fig 2 for "MIC13 and SLP2 seed the assembly of MIC60-subcomplex to facilitate crista junction formation"

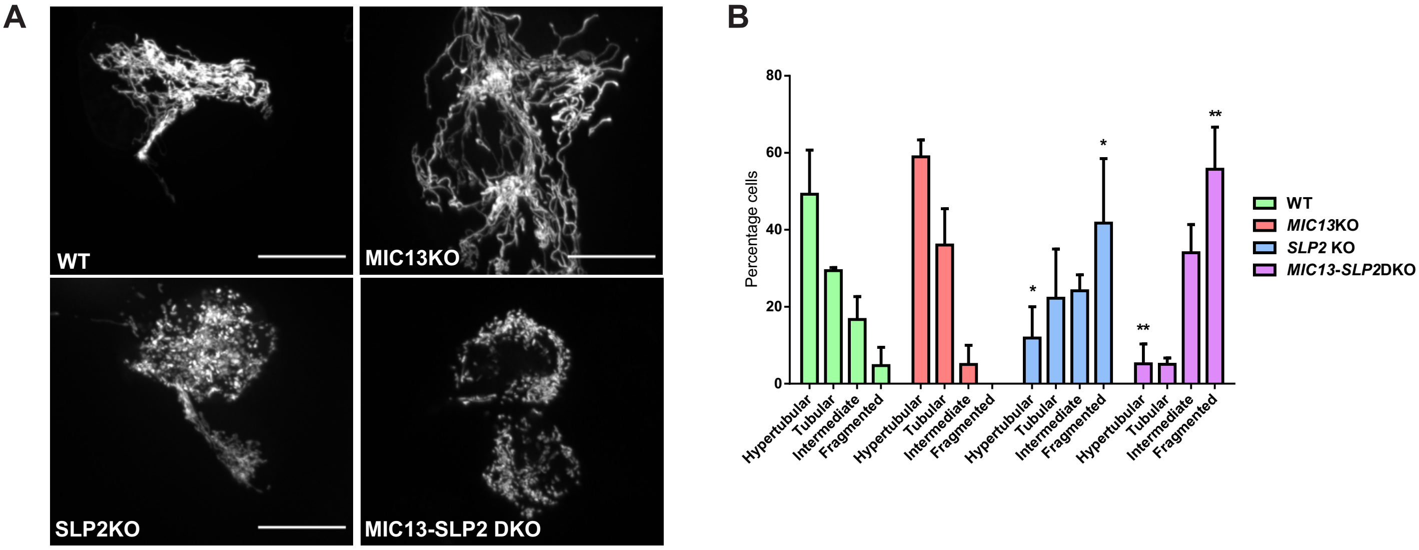


**Supplementary figure 2. SLP2 promotes stress induced mitochondrial hyperfusion independent of MICOS. A,** Assessment of mitochondrial morphologies from WT, *MIC13* KO, *SLP2* KO and *MIC13-SLP2* DKO cells post treatment with 10 μM cyclohexi mide for 2 hours. **B**, Percentage of cells displaying tubular, intermediate, fragmented or hyperfused mitochondria (n= 2). *P-value ≤ 0.05, **P-value ≤ 0.01, ***P-value ≤ 0.001. Data represented as mean with standard error of mean.Scale bar represented as 15 μm.

.
