## Supplementary Fig 3 for "MIC13 and SLP2 seed the assembly of MIC60-subcomplex to facilitate crista junction formation"

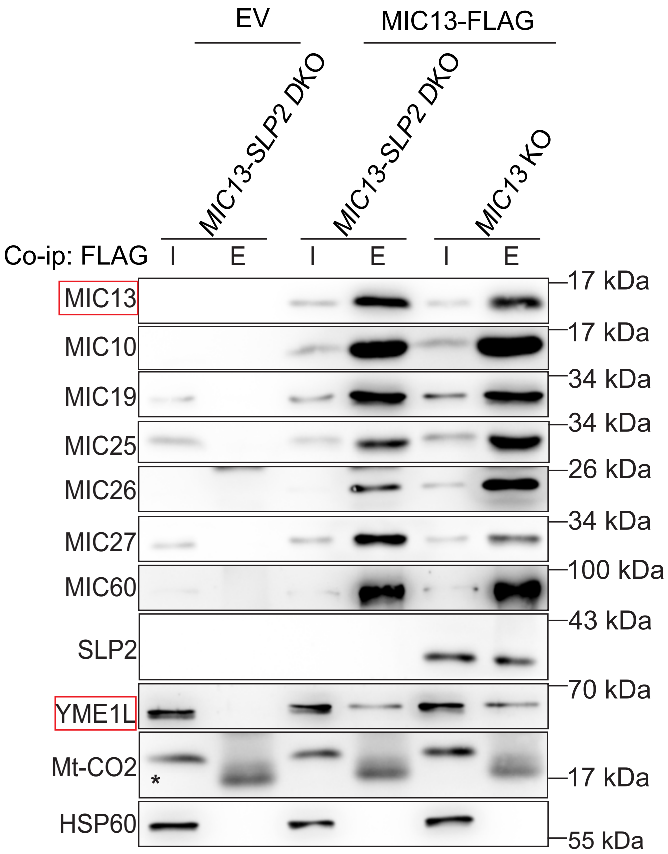


**Supplementary Figure 3.** Co-ip of MIC13-Flag using anti-Flag M2 beads with isolated mitochondria from *MIC13* KO stably expressing pMSCVpuro EV (negative control) and MIC13-Flag expressing in *MIC13* KO and *MIC13-SLP2* DKO cells. The interaction of MIC13 with other MICOS subunits was unaffected upon loss of SLP2. YME1L was found as a novel interactor of MIC13. MIC13 -YME1L interaction is independent of SLP2.
