## Supplementary Fig 4 for "MIC13 and SLP2 seed the assembly of MIC60-subcomplex to facilitate crista junction formation"

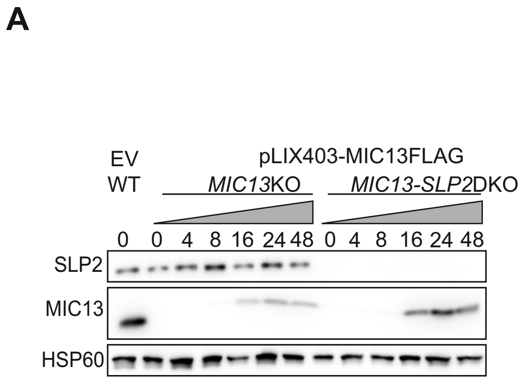


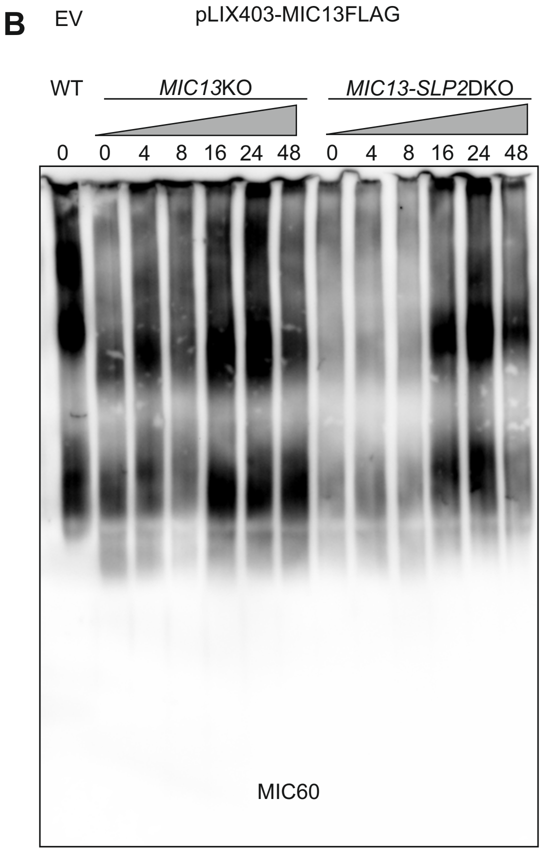


**Supplementary Figure 4. SLP2 specifically regulates assembly kinetics of MIC60. A,** WT cells stably expressing pLIX403 EV and *MIC13* KO, *MIC13-SLP2* DKO cells stably expressing pLIX403-MIC13-FLAG were treated with 1 µg/ml of doxycycline (Dox) for indicated time points. **B,** Blue native PAGE with isolated mitochondria from WT cells stably expressing pLIX403 EV, and *MIC13* KO and *MIC13-SLP2 DKO* cells stably expressing pLIX403-MIC13-FLAG treated with 1 µg/ml of Dox for indicated time points was probed for MIC60 antibody. It shows that kinetics of MIC60 assembly was dependent on SLP2.
